# Comparative metagenomics evidence distinct trends of genome evolution between sponge-dwelling bacteria and their pelagic counterparts

**DOI:** 10.1101/2020.08.31.276493

**Authors:** Joseph B. Kelly, David Carlson, Jun Siong Low, Robert W. Thacker

## Abstract

Prokaryotic associations with sponges are among the oldest host-microbiome relationships on Earth. In this study, we investigated how bacteria from several phyla have independently adapted to the sponge interior by comparing metagenome-assembled genomes of sponge-dwelling and pelagic bacteria sourced from broad phylogenetic and geographic samplings. We discovered that sponge-dwelling bacteria have more energetically expensive genomes and share patterns of depletion and enrichment for functional categories of genes that evidence evolution towards lower pathogenicity. We also identified a new defining genomic characteristic of sponge-dwelling bacteria that is virtually absent from pelagic bacteria, the presence of cassettes that contain eukaryotic steroid biosynthesis genes. Collectively, these results illuminate the trends in genome evolution that are associated with a sponge-dwelling life history strategy and have implications for furthering our understanding of how sponge-microbial symbioses have persisted through deep evolutionary time.

**Importance:** Much attention has recently been devoted to investigating the evolution of microbes that live in symbiosis with sponge hosts using microbial metagenomic data. However, several biological questions regarding this symbiosis remain unanswered. Two questions that we address here are: 1) what are the long-term consequences of the symbiosis on the evolution of microbial symbiont genome size, protein content, and nucleotide content, and 2) how is the evolution of virulence in sponge-dwelling microbial symbionts, which generally undergo a mixed transmission modes (e.g. horizontal and vertical), related to long-term stability of the symbiosis? By employing the largest comparative metagenomic analysis to date in terms of host sponge species and geographic representation, we address these questions and provide further resolution into the evolutionary processes that are involved in mediating the crosstalk between sponge hosts and their microbial symbionts.

## Introduction

The universal association of microbes with multicellular hosts is now recognized as an established principle of biology (1). In the past few decades, the combination of plummeting costs of high throughput sequencing and key statistical and computational advancements have driven a renaissance in microbiology focused on investigating the genetic underpinnings of microbial physiologies. Consequently, science has gained unprecedented insights into prokaryotic defense systems, molecular machinery related to virulence (a term that we use here to refer to both pathogenicity, the process of causing harm to the host, and the promotion of the microbe’s residence in the host), and the metabolisms of uncultured microbes. Understanding the ways in which microbes evolve in association with their hosts is central to understanding how microbes impact ourselves and our environment. In the current study, we seek to elucidate what the impacts of host-microbial associations have been on prokaryotic genome evolution in what is perhaps the oldest-standing relationship between microbes and multicellular life.

Prokaryotes are unusually successful in their colonization of sponges (phylum Porifera), a symbiosis established early in the evolutionary history of the phylum around 540 million years ago (2). Sponge microbiomes can account for up to 40% of the sponge’s volume (3) and routinely comprise thousands of microbial taxa from dozens of eubacterial and archaeal phyla (4). The taxonomic richness of sponge-dwelling prokaryotes (SDPs) is met by their physiological diversity as numerous autotrophic and heterotrophic nutritional strategies have proliferated in these microbes in addition to their capacity to synthesize a multitude of bioactive secondary metabolites (5–8). Sponge microbiomes are assembled horizontally through transmission from the environmental microbial pool and vertically through parental inheritance in a phenomenon termed mixed-mode transmission (9,10). The longevity of the SDP-sponge symbiosis is at odds with the prevalence of horizontal transmission in the system, since competition among horizontally transmitted SDPs and their relaxed dependence on survival of the host would incentivize the evolution of higher virulence, resulting in greater damage to the host that would ultimately endanger the symbiosis (11–14). The mystery stands: among the distantly related SDPs that possess such metabolic diversity, and whose transmission mode presents an evolutionary paradox, what features do they have in common that account for their shared success within the environments of sponge bodies and the stability of the symbiosis?

The current study seeks to illuminate what the consequences of sponge inhabitation have been on SDP genome evolution by performing the largest comparative metagenomic analysis to date of SDPs and free-living prokaryotes. Since the host sponge species in our analysis were sourced worldwide from multiple biomes (Table S1), we decided to use the MAGs assembled from the Tara Oceans pelagic water samples (15) as our free-living dataset to standardize the comparison. We focus on two general genomic facets; the first is the evolution of general genomic characteristics and encompasses genome size and, particularly, how it scales with protein-coding and GC contents. In the open ocean, these features are intrinsically linked to nutrient availability, which drives genome streamlining of pelagic prokaryotic genomes and A-T biases in nucleotide composition (16–19). The genomes of at least one bacterial genus, *Synechococcus* (phylum Cyanobacteria), have been found to be of comparable size between sponge-dwelling and pelagic strains (20,21). Yet, we lack a model describing how genome size scales with protein and GC content across SDPs from distantly related prokaryotic clades. Second, we investigate the shared enrichment patterns of individual genes in SDPs with the goal of elucidating the sources of selection that are experienced across SDP taxa in multiple host species. Previous work has uncovered patterns of enrichment for functional categories of genes that are characteristic of SDPs and are likely adaptations to a symbiotic lifestyle including ATP-binding cassette (ABC) transporters, prokaryotic defense mechanisms (e.g. CRISPR-CAS and restriction-modification (RM) systems), and eukaryotic-like proteins (ELPs) (22–26). Other functional categories of genes that concern virulence and are thus relevant to how the SDPs interact with their hosts, such as cellular motility and chemotaxis, have been reported as being depleted in three sulfur-oxidizing gammaproteobacterial SDPs (27) and in SDPs belonging to *Rhodospirillaceae* (Alphaproteobacteria) (28). However, theoretical work has posited that horizontally transmitted symbionts would benefit from increased motility and chemotaxis as these traits would result in increased recruitment to the host (29). Since mixed-mode transmission is a major driver of sponge microbiome assembly (9,10), we need to develop a more complete understanding of how virulence has evolved in SDPs to better model how environmental recruitment shapes the microbiomes of sponges.

## Materials and Methods

### DNA extractions and sequencing

New metagenome assembled genomes (MAGs) were produced for this study from two specimens of each of three nominal *Ircinia* species and seven *Ircinia* growth forms resulting in a sample size of 20. In Bocas del Toro, Panama, specimens of the *‘*Massive A pink’ growth form were collected from prop roots of the mangrove hammock at Inner Solarte (latitude, longitude: 9.3058, −82.1732), individuals of the growth forms ‘Massive A green’ and ‘Massive B’ from seagrass beds near the Smithsonian Tropical Research Institute’s Bocas del Toro Research Station (9.3517, −82.2590), and individuals of the ‘Encrusting’ growth form from patch reefs at Punta Caracol (9.3771, −82.3023). In Florida, specimens of *I. campana* and the ‘Ramose’ growth form were collected from a seagrass bed near the MOTE Marine Laboratory and Aquarium’s Elizabeth Moore International Center for Coral Reef Research & Restoration (24.6609, −81.4563). In Belize, specimens of the ‘BZ1’ growth form were collected from the prop roots of mangrove hammocks near the Blue Ground site (16.8083, −88.1496), specimens of *I. strobilina* and the ‘BZ2’ were collected from the coral patch reefs of Blue Ground (16.8010, −88.1461), and specimens of *I. felix* were collected from the forereef of Carrie Bow Cay (16.8042, −88.0796). Thumb-size tissue fragments were cut from each sponge and stored in 90% EtOH that was replaced at the 24 and 48 timepoints to ensure thorough inundation. Photos and descriptions of the growth forms and maps of the sampling locations can be found in Kelly et al. (2020). Note: the specimens used in the current study are present in the set of specimens used in Kelly et al. (2020).

DNA was extracted from each specimen using the Molzyme Molysis DNA isolation kit, which depletes eukaryotic host DNA in a sample resulting in an extraction that is enriched for microbial DNA. The absence of host DNA was evaluated by amplifying the Molzyme DNA isolations with the *Ircinia*-specific *cytochrome oxidase c subunit 1* primers and PCR conditions reported in Kelly & Thacker (2020). To provide a positive control, PCR amplifications were repeated for the same samples using bulk-tissue DNA isolations produced using the Wizard Genomic DNA Purification kit. Whole-metagenome shotgun sequencing was performed for the Molzyme DNA extractions after confirming the absence of an amplification band at the Yale Center for Genome Analysis on a NovaSeq6000 with a target of 30 million 2×150 reads per specimen.

### Metagenomic analysis

The reads were filtered and trimmed using FastP (32) with default parameters. Since the DNA libraries were prepared in a facility that routinely handles vertebrate model organisms, bbsplit.sh was used to remove contaminating reads mapping to the *Mus musculus* and *Homo sapiens* genomes as a precautionary step in addition to removing the control PhiX reads. Metagenomic contigs were assembled independently for each specimen using Megahit v1.2.9 (33) and subsequently binned using MetaBAT 2 v2:2.15 (34) with default parameters. Contaminating contigs were identified on the basis of being outliers with regard to genomic content (i.e., %GC and tetranucleotide content) and removed from the bins using refinem v0.0.25 (35). The qualities of the refined bins were assessed using checkm v1.1.2 (36); only bins with qualities equal to or higher than 40, calculated as genome completeness – 5x contamination (35), were retained for downstream analysis. The remaining bins were screened for uniqueness within host specimens using checkm.

To investigate the microbial mechanisms that promote their residence in sponge hosts, we compared a dataset of SDP MAGs with a dataset of previously assembled MAGs from the Tara Oceans expedition (15). The Tara Oceans dataset was produced from DNA isolated from filtered oceanic seawater, and thus we assume that most of these microbes are pelagic; these are referred to as the pelagic dataset throughout the manuscript (15). Prior to quality control and dereplication, the pelagic dataset consisted of 2307 MAGs (NCBI bioproject PRJNA391943, Table S2). The SDP MAG dataset was produced by combining our Caribbean *Ircinia* MAG dataset with 689 publicly available draft genomes of SDPs (Table S2). Prior to dereplication and further QC, the SDP dataset consisted of 1579 MAGs. The SDP and Tara Oceans datasets were independently dereplicated using drep v2.5.2 (37) with a cutoff of 96.5% identity, an estimate of maximum prokaryotic intraspecific genomic divergence backed by empirical evidence (38). We retained only MAGs with a quality equal to or greater than 40 and a completeness greater than 85% for downstream analyses. Taxonomy was again assigned and a whole-genome phylogenetic tree was produced for the combined datasets using GTDB-Tk v1.0.2 (39).

Protein predictions and subsequent KO annotations (http://www.kegg.jp/kegg/) were performed on the MAGs using Enrichm v0.5.0 (40). Domain PFAM annotations were performed on the MAG proteins using Interproscan v5.39-77 (41). For proteins that contained duplicated identical domain annotations, only one domain was retained so as to avoid double counting proteins.

### Statistical analysis

To identify features that were significantly enriched in one life history category relative to the other, we performed pairwise permutational t-tests, each run from 10,000 iterations, comparing the average abundance (e.g. copy number) of a feature in the pelagic vs. the SDP dataset. To factor for differences in genome size, we normalized feature counts by dividing them by the total number of predicted proteins prior to the t-tests. P-values were corrected using the FDR adjustment of Benjamini-Hochberg (42) using the total number of features within an annotation category (either KO or PFAM). Given the high number of multiple comparisons, we further decided to assign a conservative significance cutoff of p<0.01. Additionally, we omitted features that were present in less than 10% of both the SPD and pelagic MAGs.

Linear models (linear regressions) were inferred to describe the relationship between genome size vs. GC content and coding density. Statistical significance in differences between the linear models of SDP and pelagic microbes was inferred using ANCOVA. We tested for significance in genome size, GC content, coding density, and the total number of proteins between SDPs and pelagic MAGs within a phylogenetic framework using an ultrametric bacterial tree estimated with the software PATHd8 (43). The GTDB-Tk bacterial phylogeny was used as the input and node dating was calibrated using the divergence times estimates of Battistuzzi et al. (2004) for the cyanobacteria/actinobacteria, cyanobacteria/proteobacteria, and alphaproteobacteria/gammaproteobacteria splits. We then performed phylogenetic ANOVAs for each of the aforementioned genomic traits using the R package geiger (45), each for 10,000 iterations. All aforementioned tests of significance and regressions were performed in R v.3.6.0.

## Results and Discussion

To investigate shared patterns of genome evolution in SDPs, we performed a comparative metagenomic analysis evaluating nucleotide composition, genome size, and gene content of SDPs from a broad sampling of SDP MAGs sourced from multiple host species and oceanic basins against the MAG dataset of pelagic prokaryotes sequenced during the Tara Oceans project (15). After screening for quality, completeness, and dereplication, our final bacterial dataset was composed of 479 SDP MAGs sourced from 23 nominal and 10 undescribed host species and 629 pelagic MAGs; representatives from the Atlantic, Mediterranean, Red Sea, and Pacific Ocean basins were present in both datasets (Table S1). Twenty-five bacterial phyla were recovered with a high degree of phylogenetic overlap between the two datasets, with the majority of SDP MAGs (95.0%) and pelagic MAGs (92.7%) belonging to phyla that contained both life history strategies (Table S4). We report our results below for analyses performed on the bacterial MAGs, as only 15 archaeal MAGs were recovered (Table S5). We confirmed that the omission of Archaea did not change the interpretations of the results by repeating our analyses on the total (Bacteria + Archaea) MAG dataset (data not shown). We tested the hypotheses that 1) general genomic characteristics distinguish SDPs from free-living oceanic bacteria and 2) enrichment patterns in functional categories of genes that are relevant to host-symbiont crosstalk, including genes that facilitate horizontal transmission (e.g. motility and chemotaxis), are different in SDPs relative to pelagic bacteria.

### SDPs have evolved larger genomes with higher GC contents

The evolution of general genomic features across marine prokaryotic taxa has been shaped by life history strategies. We found that SDPs have larger genomes, with both more proteins and non-coding DNA, and higher GC content (Fig. 1, 2). The effect of life history strategy remained significant after accounting for phylogenetic relatedness (Fig. 1). Life history strategy also had a significant effect on the relationship between genome size and both GC content (ANCOVA: F_1,1104_ = 118.32, p < 2e^−16^) and coding density (ANCOVA: F_1,1104_ = 104.3, p < 2e^−16^) (Fig. 2). Based on comparisons among regression coefficients, we found that genome size was a stronger predictor for GC content and coding density in pelagic MAGs relative to SDPs (Fig. 2). Our finding of larger genome sizes and higher GC contents in SDPs relative to seawater bacteria is consistent with previous observations made of SDPs inhabiting three Mediterranean sponge species, *Petrosia ficiformis*, *Sarcotragus foetidus*, and *Aplysina aerophoba*, suggesting that these are geographically widespread genomic characteristics found in SDPs across host taxa (24).

**Figure 1.**
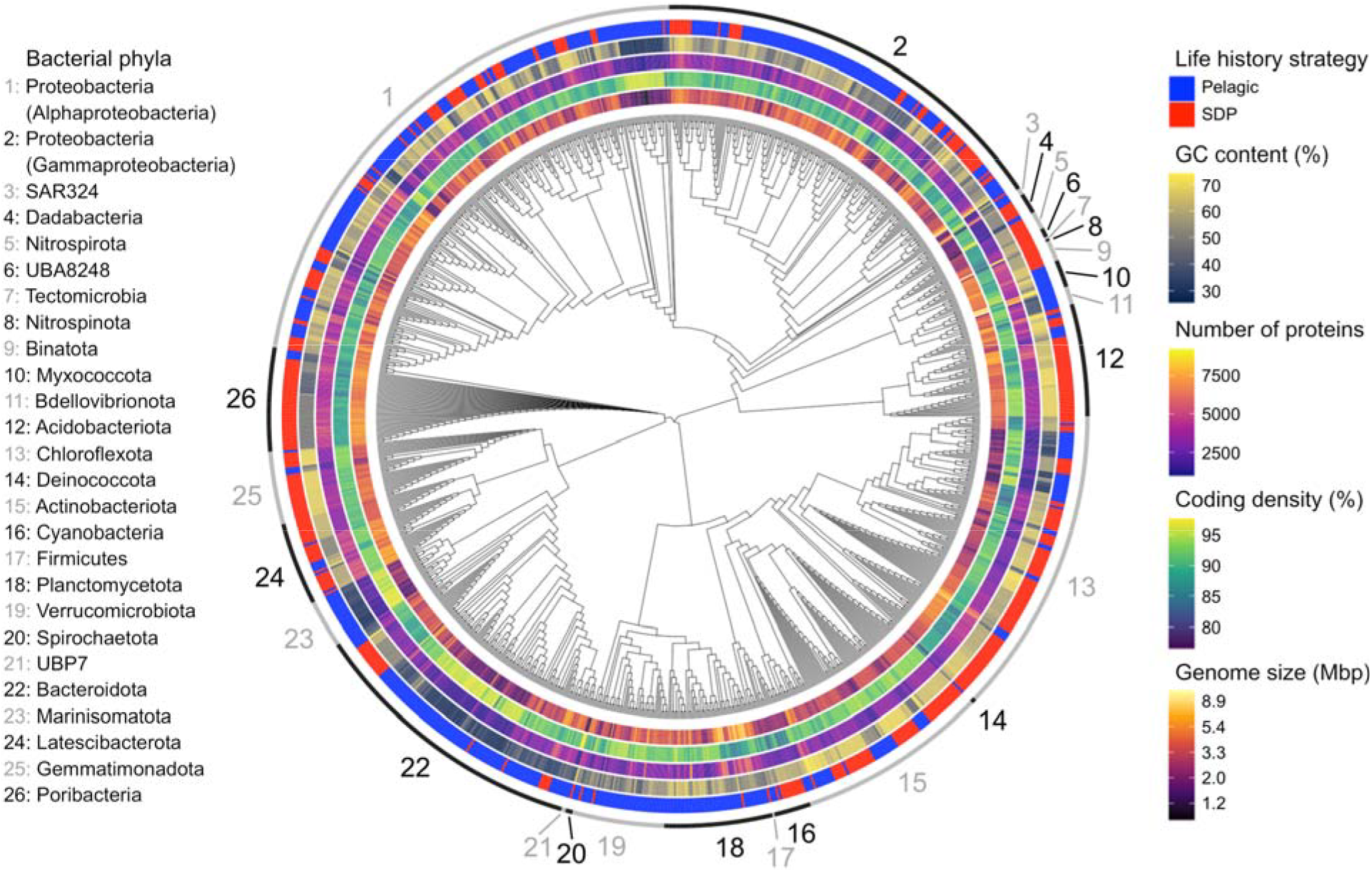
The phylogenetic distribution of endogenous genomic traits in bacteria is independent of life history strategy. Phylogenetic ANOVAs of genomic characteristics show that they are significantly different between SDPs and pelagic MAGs after correcting for phylogenetic relatedness and multiple comparisons (fdr): genome size (F_1,1106_=68.218, p=0.0488), coding density (F_1,1106_=188.62, p=0.0005), number of proteins (F_1,1106_=66.227, p=0.0488), GC-content (F_1,1106_=292.44, p=0.00025). Bacteria phyla are annotated by the alternating gray and black sections of the outer circle.

**Figure 2.**
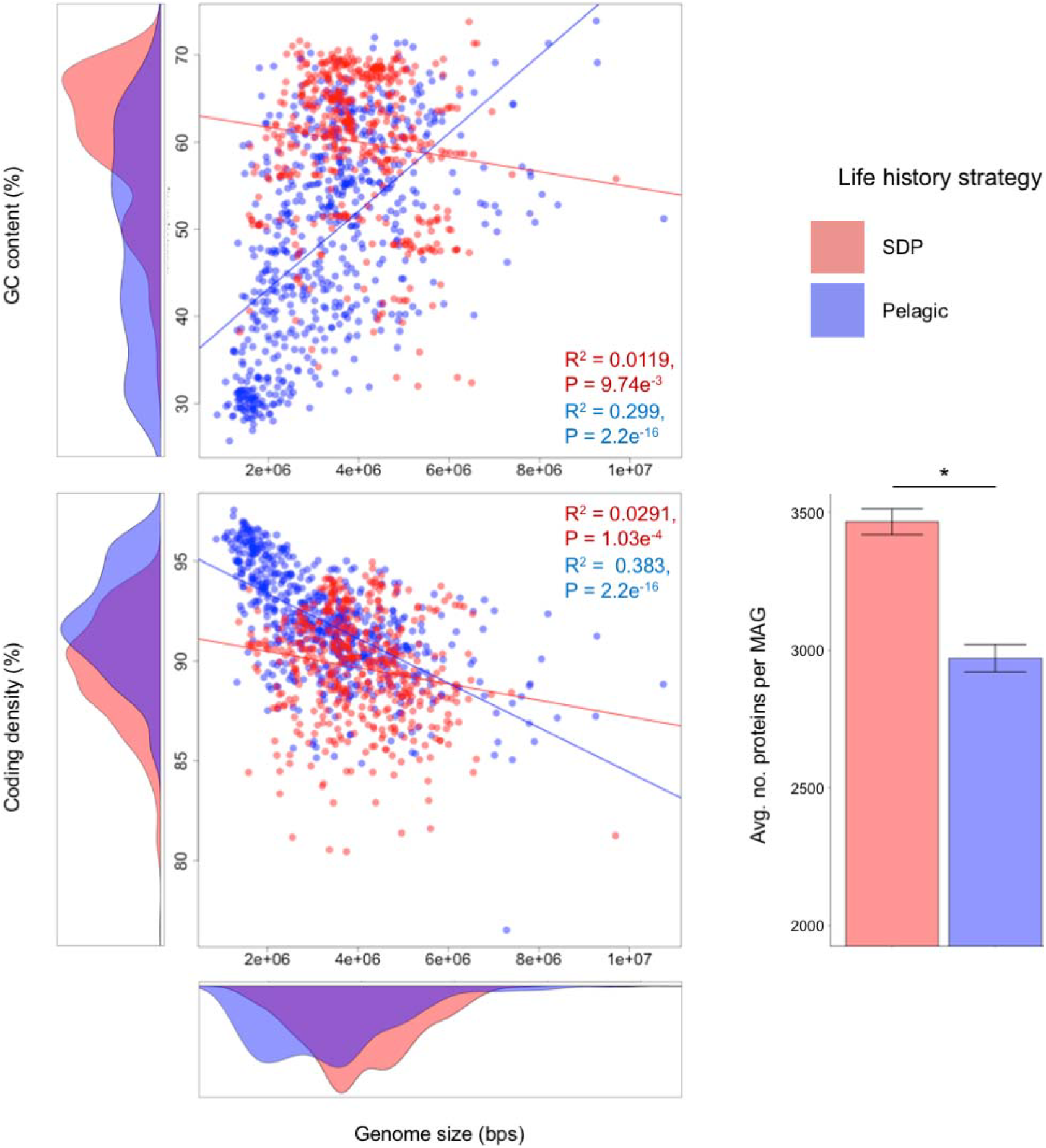
Genomes of SDPs are more energetically expensive than pelagic genomes. **A:** Scatterplot comparing relationships prokaryotic genome sizes to GC content and **B**, genome size to coding density for SDPs and pelagic microbes. Coding density is reported as the percentage of basepairs that are included in a coding region. R^2^ and statistical significance is reported for linear models inferred for the pelagic and SDP MAGs. Plots to the left of the y-axes and below the x-axes are kernel density estimation plots. Degrees of freedom for SDP MAGs = 477, pelagic MAGs = 627. **C:** Bar chart comparing average number of proteins predicted from SDP and pelagic MAGs (phylogenetic ANOVA reported in the caption of Figure 1). Asterisks here and elsewhere in figures denote significance.

We hypothesize that differences in energy availability between the two life history strategies may account for the contrasting genomic characteristics of SDP and pelagic MAGs. Due to nitrogen limitation in the open ocean, pelagic prokaryotes tend to have streamlined genomes and lower GC content; this strategy reduces the energetic costs of DNA replication (16–19). Each of these expectations are met in our pelagic dataset, as pelagic MAGs have smaller genomes with fewer genes, less non-coding DNA, and lower GC content. However, opposite trends in genome content were discovered in SDPs, suggesting that the sponge host might be providing the symbionts with nitrogen. This hypothesis is corroborated by the observation that two sponge species, *Xestospongia muta* (46) and *Haliclona cymiformis* (47), are documented sources of inorganic nitrogen (NH_3_) for SDPs. Further research could investigate whether the exploitation of nitrogenous waste by SDPs is widespread throughout sponge host taxa to elucidate whether a sponge-dwelling lifestyle alleviates SDPs of nitrogen limitation and enables the maintenance of larger genomes with higher GC contents.

### Gene enrichment patterns suggest SDPs are equipped to exploit the organic molecules present in the sponge body and have evolved towards lower pathogenicity

Previous studies have concluded that several classes of molecular transporters (e.g. ABC transporters) are enriched in SDPs (22,25,48). Our results are consistent with this finding (Fig. 3) and provide further resolution into which classes of ABC transporters are enriched or depleted in SDPs. For example, SDPs are depleted for lipopolysaccharide (LPS) transporters, Hg^+^/heavy metal efflux proteins, and the *pha* family of pH-dependent H^+^ cation transporters. Conversely, and consistent with observations made in previous studies (22,28,48–50), SDPs are enriched for amino acid and spermidine/putrescine transporters. Early work discovered that concentrations of free amino acids are relatively high in sponges, which are thought to play a role in host osmoregulation (51). Microbial symbionts could be capitalizing on this pool of bioavailable nutrients to supplement their own metabolism, possibly to provide cheap sources of amino acids for protein biosynthesis. Likewise, spermidine and putrescine are polyamines that are essential for cell growth (52) and perform protective functions for host-associated prokaryotes (53). Recent genomic analysis evidenced a capacity for members of at least one clade, the gammaproteobacterial order Tethybacterales, to metabolize and derive energy from spermidine (54). Collectively, our results suggest that SDPs have evolved a capacity to utilize the organic molecules in sponge bodies to meet their energetic needs.

**Figure 3.**
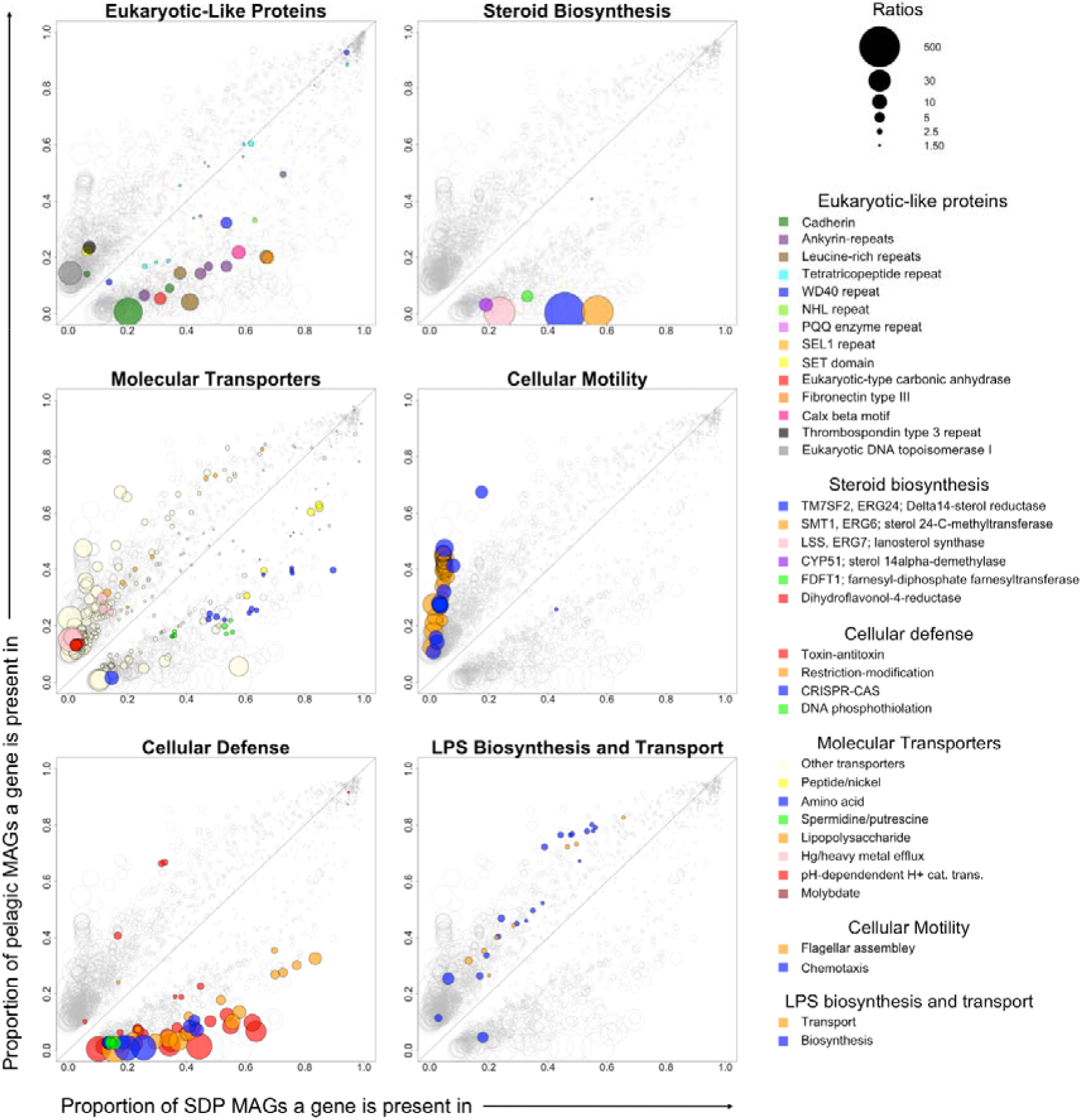
SDPs are well-defended against foreign DNA, are equipped to promote their residence in host sponges, and have reduced pathogenicity. Plots depicting genes that are enriched in pelagic (i.e. depleted in SDPs) or enriched in SDPs. Each circle corresponds to a gene that was determined as being differentially abundant in either SDPs or pelagic MAGs in terms of normalized average gene copy number (see methods). The size of each circle is scaled to the log-transformed ratio of the gene calculated as (avg. copy number SDP) / (avg. copy number pelagic) for genes under the y=x diagonal and (avg. copy number pelagic) / (avg. copy number SDP) for genes above the y=x diagonal. Non-transformed ratios are reported in the legend. The ELP graph was constructed from the Interproscan annotations; all others were constructed using the KO annotations produced from enrichm.

Consistent with the observations made for gammaproteobacterial SDPs made by Tian et al. (2017) and Karimi et al. (2018), we detected a near complete absence of motility and chemotaxis genes across SDPs. Additionally, we detected a depletion of lipopolysaccharide (LPS) biosynthesis genes and the aforementioned LPS transport genes in SDPs (Fig. 3). We also recovered enrichment patterns of ELPs including the previously reported fibronectin type-III, ANK, LRR, TPR, NHL, WD40, cadherin, and PQQ domains (22,25,26,48,55) and two new ELPs: calx-beta motifs and eukaryotic-type carbonic anhydrase. Most notably, we discovered that SDPs were also enriched for genes involved in steroid biosynthesis that are localized on the bacterial chromosomes in cassettes (Fig. 4) that constitute an incomplete eukaryotic steroid biosynthesis pathway (KEGG pathway ko00100). Other genes present in this pathway were found to be missing from the SDP genomes. Our results are similar to prior observations made of SDPs inhabiting the sponge *Vaceletia sp*. where, again, these genes represented an incomplete ko00100 pathway that lacked the machinery to produce steroid precursor molecules. Instead, it was found that several complementary components of the pathways were discovered in the host *Vaceletia* genome, suggesting that steroid precursors may originate in the sponge itself (56).

**Figure 4.**
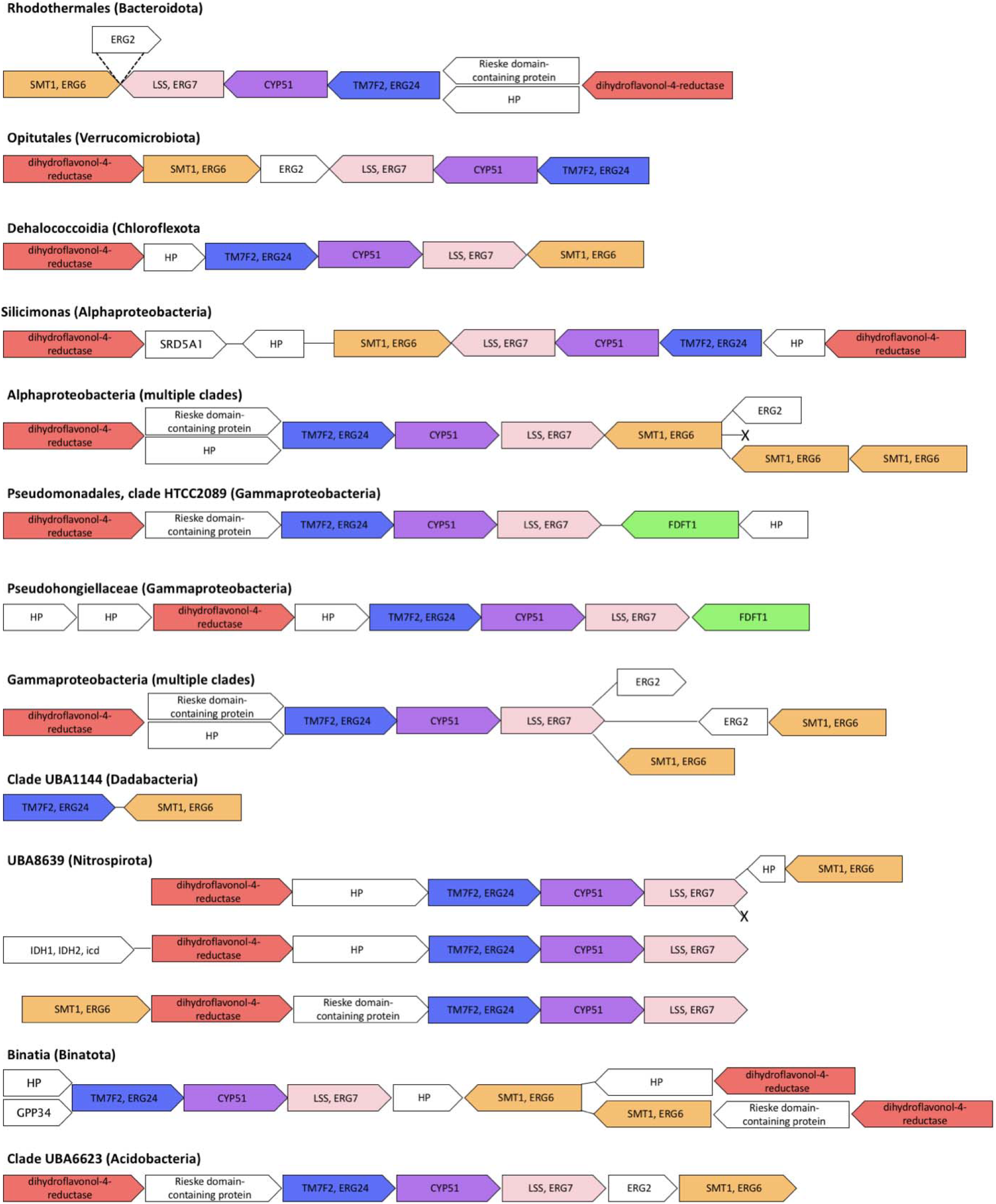
SDPs contain partial eukaryotic steroid biosynthesis pathways. Representative cassettes of steroid biosynthesis genes are depicted. Genes in parallel denote common alternative combinations for a clade. TM7SF2, ERG24, Delta14-sterol reductase [EC:1.3.1.70] (K00222); SMT1, ERG6, sterol 24-C-methyltransferase [EC:2.1.1.41] (K00559); LSS, ERG7, lanosterol synthase [EC:5.4.99.7] (K01852); CYP51, sterol 14alpha-demethylase [EC:1.14.14.154 1.14.15.36] (K05917); FDFT1, farnesyl-diphosphate farnesyltransferase [EC:2.5.1.21] (K00801); and ERG2, C-8 sterol isomerase [EC:5.-.-.-] (K09829) are components of the steroid biosynthesis pathway (ko00100). Dihydroflavonol-4-reductase [EC:1.1.1.219] (K00091), an oxidoreductase acting on the CH-OH group of donors with NAD+ or NADP+ as acceptor, is not assigned to a pathway and was included here and in Figure 2 due to its consistent association with genes belonging to the steroid biosynthesis (ko00100) pathway and its likely relevant biochemical activity. ‘HP’ abbreviates hypothetical protein. Arrows denote direction of the coding sequence. Genes are colored to match the legend of the ‘Steroid Biosynthesis’ plot of Figure 2. ERG2 is uncolored as it was removed from the data producing Figure 2 due to being in low proportions (<10%) of both the SDP and pelagic MAG datasets. Note: genes are not drawn to scale by length.

Proteins involved in motility, such as flagellin, and bacterial lipopolysaccharides are directly involved in eliciting the immune response from the host that causes pathogenicity (57–60); thus, a potential evolutionary outcome of the depletion of these genes in SDPs might be one of reduced symbiont pathogenicity. At face value, this hypothesis appears to be at odds with the theory that lower levels of pathogenicity would only be expected to confer a selective advantage to the microbe if the persistence of its genome was completely dependent on survival of the host, i.e. under obligate vertical transmission (11,12). On the other hand, higher rates of pathogenicity are expected to evolve in microbes exhibiting mixed-transmission modes as pathogenic mechanisms act to protect the microbe, for example through the strengthening of cell envelopes with LPS, while not damaging the microbial population’s growth rate since it is not linked to the host’s (13). Why then do we detect a depletion of pathogenic mechanisms in prokaryotes that are likely undergoing mixed-modes of transmission in sponges (9,10)? One possibility is that the products of the steroid biosynthesis gene cassettes could be playing an analogous role to LPS in strengthening the cellular envelope and contributing to the resistance of stressors such as antimicrobial secondary metabolites (5,61) and hypoxic conditions (62,63) that are present in the interior of the host, as is the case in other symbiotic microbes such as the legume-inhabiting *Bradyrhizobium* (64–66). An alternative role for the steroid genes might be that their products have immunosuppressive properties, similar to the endogenous steroid hormones of eukaryotes, which could help prevent removal of the SDPs by the hosts’ immune system (67,68). Under this scenario, SDPs could still experience selection against LPS, cellular motility, and chemotaxis genes if the dependence of SDPs on their hosts for survival outweighs the fitness benefits that these genes confer outside of the host. Experimental determination of the gene products of the cassettes containing steroid biosynthesis genes and functional assays are required to delineate the roles that the steroid genes play in the sponge-SDP symbiosis.

## Conclusion

Our results provide new insights into the characteristics that distinguish SDP from those of free-living microbes and help elucidate the metabolic and defensive strategies adopted by SDPs that translate to their success in sponge hosts. We discover greater genome sizes and GC contents of SDPs could that could, in part, be enabled by allocation of nitrogen from the host sponge to SDPs via nitrogenous waste products. Additionally, we discovered the near complete absence of genes related to cellular motility chemotaxis and a depletion of LPS biosynthesis and transport genes in SDPs, evidence that inhabitation of the sponge interior involves an evolution towards decreased pathogenicity. Finally, we discovered a novel defining characteristic of SDPs: the presence of steroid gene cassettes. In particular, we advocate for future research to focus on identifying the products of these steroid-like pathways and identifying the roles that they play in mediating the symbiosis between sponges and SDPs.

## Acknowledgements

We thank Dr. Jackie L. Collier and Dr. Liliana Davalos-Alvarez for their input and comments on the manuscript; the staffs of the Smithsonian Tropical Research Institute’s Bocas Research Station, the Smithsonian’s Carrie Bow Cay Field Station, and the MOTE Marine Laboratory and Aquarium’s Elizabeth Moore International Center for Coral Reef Research & Restoration for their support on logistical aspects of the field work; and Barrett Brooks (Smithsonian Museum of Natural History) and Karen Koltes (form. Office of the Secretary of the Interior) for help with specimen collections in Carrie Bow Cay. We thank the Yale Center for Genome Analysis for their attentiveness in the metagenomic DNA library preparation and sequencing; the staff of the SeaWulf HPC at SBU, which we used to perform the majority of our analyses; and our funders at Experiement.com.

## Author contributions

study conceptualization, methodology: JBK, JSL, DC; formal analysis and investigation: JBK; resources and funding acquisition: JBK and RWT; writing (original draft preparation): JBK; writing (review and editing): JBK, JSL, DC, RWT; supervision: RWT.

## Data and materials availability

Raw data that was produced during this study are deposited under GenBank Accession XXX. Assembled MAGs are deposited under NCBI BioProject XXX. Accessions for publicly available data are announced in the main text and supplement.

## List of Supplementary Materials

Tables S1-S7

